# The effect of blue-blocking lenses on photostress recovery times for low and high contrast chromatic and achromatic stimuli

**DOI:** 10.1101/745000

**Authors:** Hind Saeed Alzahrani, Sieu K. Khuu, Adiba Ali, Maitreyee Roy

## Abstract

The selective reduction in visible wavelengths transmitted through commercially available blue-blocking lenses (BBLs) is known to influence the appearance and contrast detection of objects, particularly at low light levels which may impact the human retinal receptor response time to dynamic light changes during phostress events. In the present study, we assessed whether BBLs selectively affect photostress recovery times (PSRTs) in 12 participants for chromatic and achromatic stimuli presented under low and high contrast luminance conditions. Four types of commercially available BBLs were evaluated, and their effects on PSRTs were investigated. Our results showed that PSRTs required to detect high contrast chromatic and achromatic stimuli were unaffected by BBLs when compared to a clear control lens. However, PSRTs were significantly affected by BBLs and were longer when chromatic and achromatic stimuli were of low contrast. In addition, BBLs had the greatest impact on the PSRTs of blue coloured targets, and this was dependent on the spectral transmittance profile. These results indicate that wearing BBLs under low contrast conditions can have serious implications for visual behavior, particularly under low-light levels and in situations in which the observer is directly exposed to bright light sources. For example, during night time driving, the driver might be briefly exposed to bright lights by glancing at the headlights of a passing car. This increases the time required for vision to be restored after bright light exposure, resulting in delayed object detection, and therefore stoppage and reaction times, which might pose a safety risk for a driver.

## Introduction

Blue-blocking lenses (BBLs), particularly so-called yellow BBLs with cutoff wavelengths between 450 nm and 512 nm, have been designed to provide protection against hazardous blue light [1]. These yellow BBLs have become popular in recent years and are used to aid vision in tasks such as shooting [2], skiing, aviation [3], hunting and sailing [4]. It has been suggested that BBLs benefit vision with reported improvement in visual tasks such as visual acuity [5, 6] particularly enhancing the clarity of vision [7], and decreased glare [5, 8]. However, at present, the benefit of BBLs have yet to be fully verified by empirical and independent evidence [9].

Despite the potential benefits of BBLs, previous studies have shown that BBLs may impair vision, particularly affecting the ability of the visual system to detect contrast under different visual conditions [5, 10–12]. For example, Thomas et al. have shown that while BBLs do not affect sensitivity to high-contrast photopic stimuli, detection of low-contrast mesopic stimuli is impaired by BBLs [5]. BBLs also dramatically reduce contrast sensitivity under scotopic conditions even after dark adaptation [10]. Collectively, these studies suggest that BBLs have the potential to affect the visibility of targets at low lighting levels, which poses a risk for visual behaviors such as driving and visual search under twilight and night-time conditions. However, the full extent to which BBLs, particularly newer generation lenses, affect visual perception, remains unclear despite their commercial availability and prescription by optometrists [9, 13].

Previous studies have begun to address this paucity in knowledge by investigating how newer generation BBLs might affect a number of visual judgements such as colour, contrast sensitivity, and visual acuity [13, 14]. Of particular note are studies that investigated whether BBLs (typically as yellow intraocular lenses (IOLs)) contribute to the time required to recover from a brief exposure to an intense light source, in the so-called photostress test [11, 12]. Here, retinal receptors are initially driven to the maximum response by exposure to intense light, and the time required for vision to be restored (i.e., recovery time) provides an indication of photoreceptors to respond to dynamic light change and recover function [15,16–19]. The speed of recovery from a photostress event is dependent on a variety of factors such as the observer’s current adaptive state and macular pigments [19, 20]. Quantifying PSRT is particularly important for vision under twilight and night conditions, in which the visual system might be exposed to a bright light source (e.g., passing car lights) and become temporarily ‘blind’ until recovery occurs.

Two studies have investigated BBLs as a contributing factor to PSRTs [11, 12]. Hammond et al. [11, 12], quantified PSRTs to a monochromatic (yellow) sinusoidal grating in patients with yellow intraocular lenses (IOLs). However, they found that PSRTs were not significantly different from clear IOLs or phakic controls, but in a subsequent study, the authors did show that PSRTs improved when the stimulus background was blue. This improvement in PSRTs might be attributed to the fact that BBLs inherently reduce blue light, and thereby enhancing image contrast. While these reports implicate that BBLs have the potential of affecting vision and PSRTs, the stimulus conditions under which they impair vision remains unclear. In the present study, we sought to further contribute to understanding the potential effects of BBLs on PSRTs by investigating how they might be affected by stimulus contrast and colour. Our motivation for doing so is two-fold: Firstly, as mentioned, BBLs appear to greatly affect vision under low light level conditions, and they might also affect PSRTs. Secondly, BBLs selectively filter blue light, and potentially PSRTs are dependent on the colour of the stimulus [11, 12].

In the present study, two experiments were conducted to investigate the effect of newer generation BBLs on PSRTs. In Experiment 1, the time required to correctly identify an achromatic letter optotype after exposure to an intense light source was measured for low and high stimulus contrasts sufficiently for those to be within photopic and mesopic limits. In Experiment 2, PSRTs were measured for chromatic (blue, red, yellow and green) high and low contrast stimuli. For Experiments 1 and 2, PSRTs were measured using 4 commercially available BBLs (UV++Blue Control, Crizal Prevencia, Blue Guardian, and Blu-OLP lenses) and a clear lens as a control. The newer generation BBLs showed higher spectral transmission properties than previously used yellow coloured BBLs [21]. Recent studies showed the effect of commercially available BBLs on visual and non-visual functions and reported that BBLs might have significant unintended effects on visual behavior as they attenuate blue light required for blue perception and scotopic vision [21, 22]. This may pose a risk regarding their use under low lighting conditions such as night driving. However, their potential effects on visual perception have not been fully quantified in the empirical and clinical studies [9, 13, 14, 22].

## Materials and Methods

### Blue-blocking lenses (BBLs) characteristics

As mentioned, a clear control lens and 4 BBLs were utilised in the present study: UV++Blue Control (JuzVision), Crizal Prevencia (Essilor), Blue Guardian (Opticare), and Blu-OLP (GenOp). The spectral transmittance characteristics of these lenses were previously measured using a Cary 5000 UV-Vis-NIR with an integrating sphere spectrophotometer (Model: EL04043683) across a range of wavelengths from 280 to 780 nm. The outcomes of this analysis are shown in Fig 1, which demonstrate that the BBLs utilized in the present study were effective in reducing transmittance of short wavelengths of light (in comparison to the clear control lens), but the extent is dependent on the type of BBLs, with in particular the Blu-OLP and Crizal Prevencia lenses filter the most light [21, 22].

**Fig 1.**
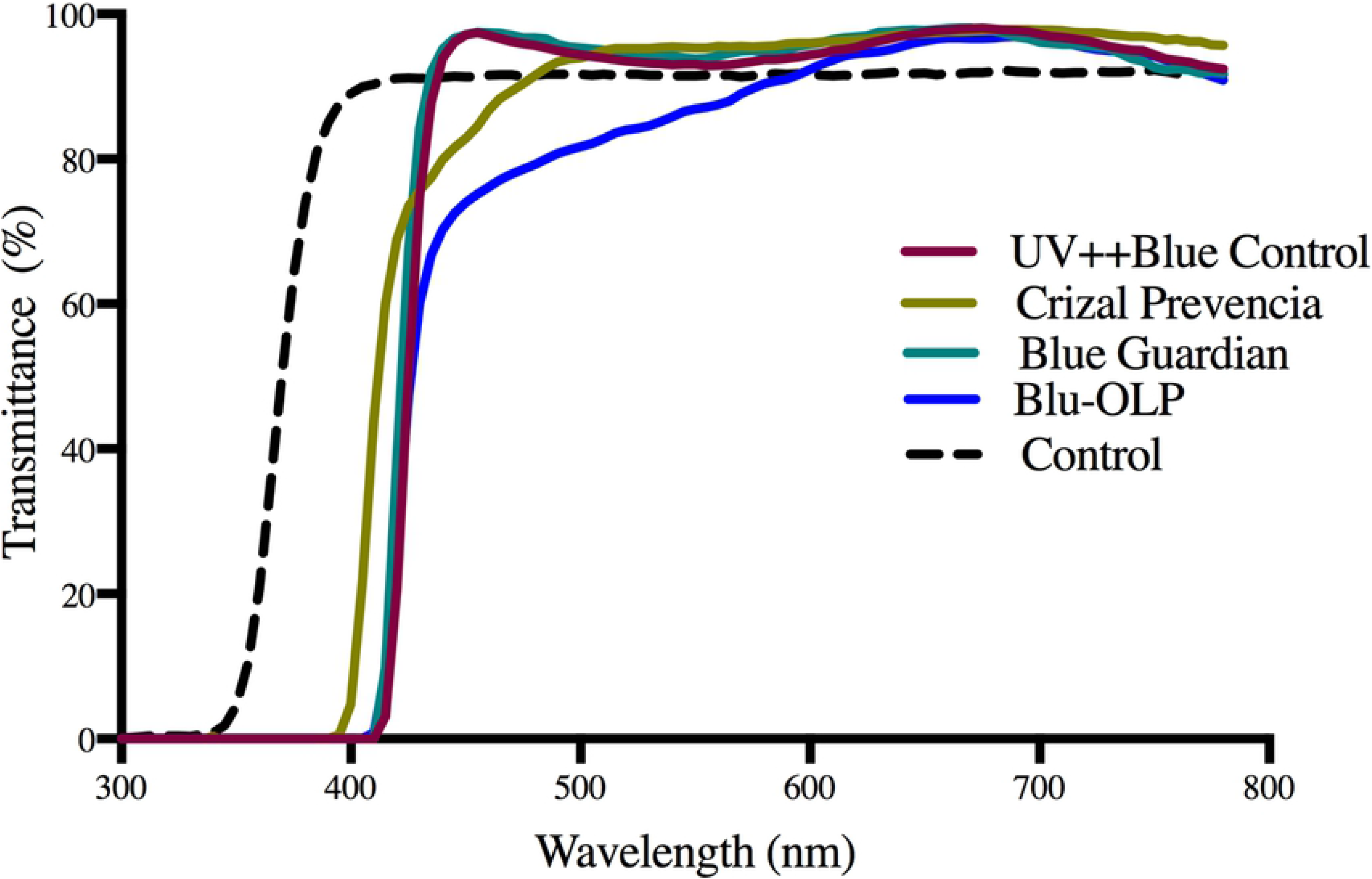
Transmission characteristics of the 4 BBLs and a clear lens as a function of wavelength.

The CIE (1931) x, y chromaticity coordinates and the luminance Y of BBLs were previously calculated with a simulated D65 illuminant. The BBLs had x, y, Y coordinates of 0.318, 0.338, 93.82 (UV++Blue Control); 0.331, 0.354, 92.20 (Crizal Prevencia); 0.320, 0.338, 95,39 (Blue Guardian); and 0.336, 0.355, 87.50 (Blu-OLP). However, these BBLs block only a small part of the range of blue wavelengths (< 420 nm), while transmitting other longer wavelengths of visible light approximately similar rates [21, 22]. All tested lenses were prepared to be worn as goggles, which allowed the BBLs to be worn over spectacles.

### Participants

Twelve participants aged between 18 and 39 years participated in the study, and they were randomly divided into two groups and participated in either Experiment 1 (n = 7) or Experiment 2 (n=5). All had normal or corrected to normal visual acuity with no history of any visual abnormalities. All participants were screened for monocular and binocular visual acuity of 6/6 or better using a Snellen chart, and normal colour perception using the Ishihara Test Book 24 Plate abridged edition. Individuals with colour deficiencies or with a history of ocular disease were excluded from this study. Each participant gave their written consent prior to testing, and the risks and benefits of the study were explained. The research adhered to the Tenets of the Declaration of Helsinki. Ethics approval was granted by the University of New South Wales Australia Human Research Ethics Advisory Panel (reference number: HC16934).

### Experiment 1: The dependency of recovery times on stimulus contrast under photopic conditions

#### Stimuli

The visual stimuli (achromatic and chromatic) used in Experiment 1 and Experiment 2 were generated on a 15-inch MacBook Pro using custom software written in MATLAB (version 14) and displayed on a linearised CRT monitor screen. The detected stimulus was a single uppercase letter Snellen optotype (2 degrees in visual angle, including: D, E, F, H, N, P, R, U, V, Z) of different Weber contrasts (as mentioned below) which was viewed on a black background (3 cd/m^2^) from a viewing distance of 130 cm, see Fig 2 as an example.

**Fig 2.**
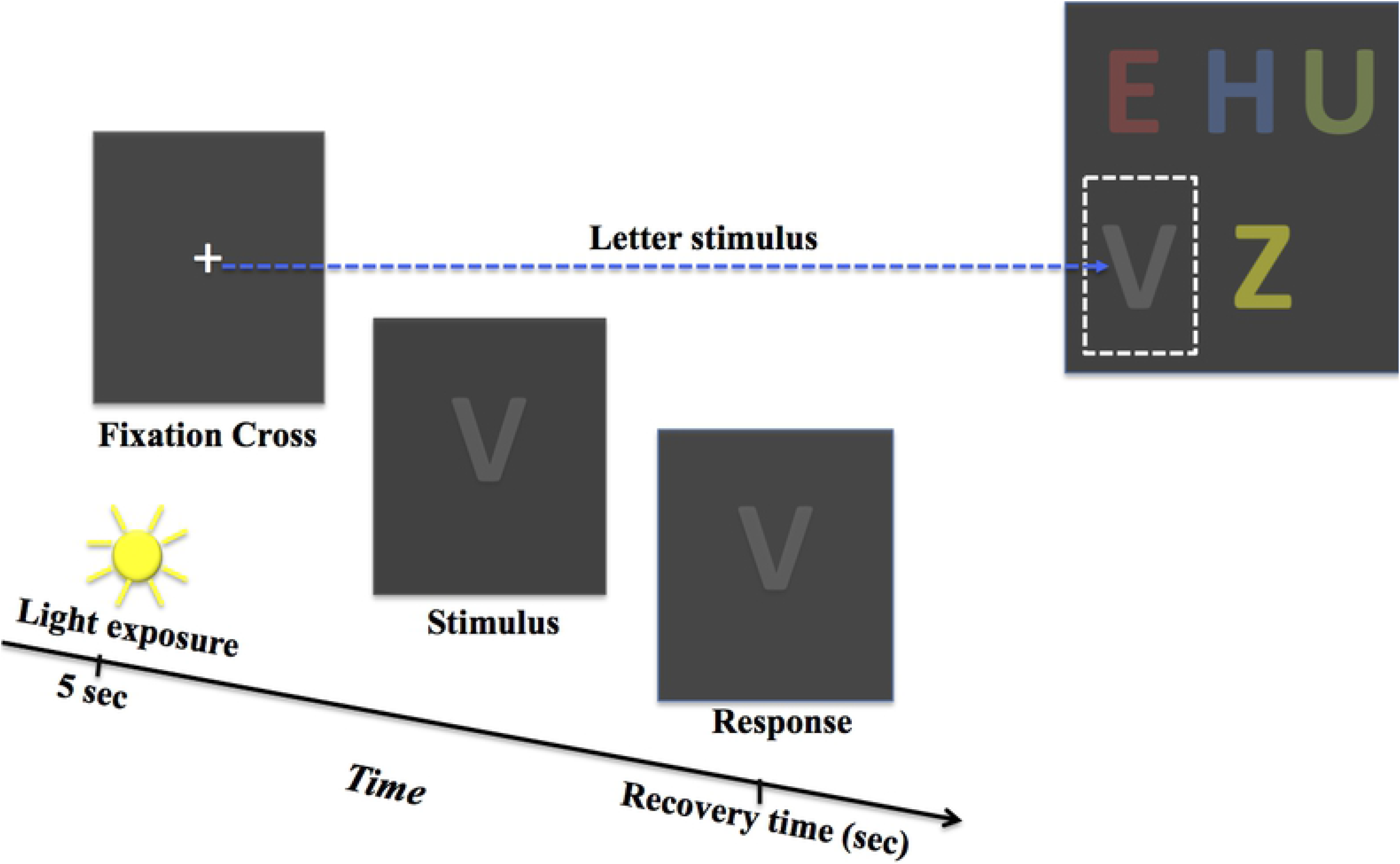
A schematic representation of a PSRT experimental trial showing achromatic and chromatic stimuli.

In Experiment 1, achromatic letter stimuli were grey and displayed on a darker grey background at Weber contrasts of 0.1, 0.2, and 0.4. Monochromatic letter stimuli were also displayed on a dark gray background and were either red (CIE 1931 xy: 0.64, 0.35), green (CIE 1931 xy: 0.3, 0.6), yellow (CIE 1931 xy: 0.51, 0.42), or blue stimuli (CIE 1931 xy: 0.15, 0.06) and corresponded to the maximum output of the sRGB monitor. All chromatic stimuli were presented at one contrast level of 0.4.

#### Procedure

In Experiment 1, the participant wore a custom-made goggle with removable lenses and was seated in front of a monitor screen. The participant viewed the monitor binocularly, and steady viewing was maintained by using a head and chin rest. The participant was allowed to adapt to wearing each google for 2 minutes before starting the experiment. The viewing distance was 130 cm from the monitor screen and was level with the participant’s eyes. Before viewing a stimulus, the participant was exposed for 5 seconds to very high light intensity from a 30W- white LED lamp (5000K, 12640lux, 28727cd/m^2^) mounted at a distance 30 cm from the eyes along the fixation axis, and under safe conditions [23]. The high-level of light from the LED lamp is sufficient for cone photoreceptors to respond and reach maximum response, as shown in Fig 3 which describes the relative sensitivity of cone photoreceptors in response to the light source. After photostress, the optotype was immediately presented at the centre of the computer monitor, and the time required to correctly name the letter provided an indication of the PSRT (Fig 2).

**Fig 3.**
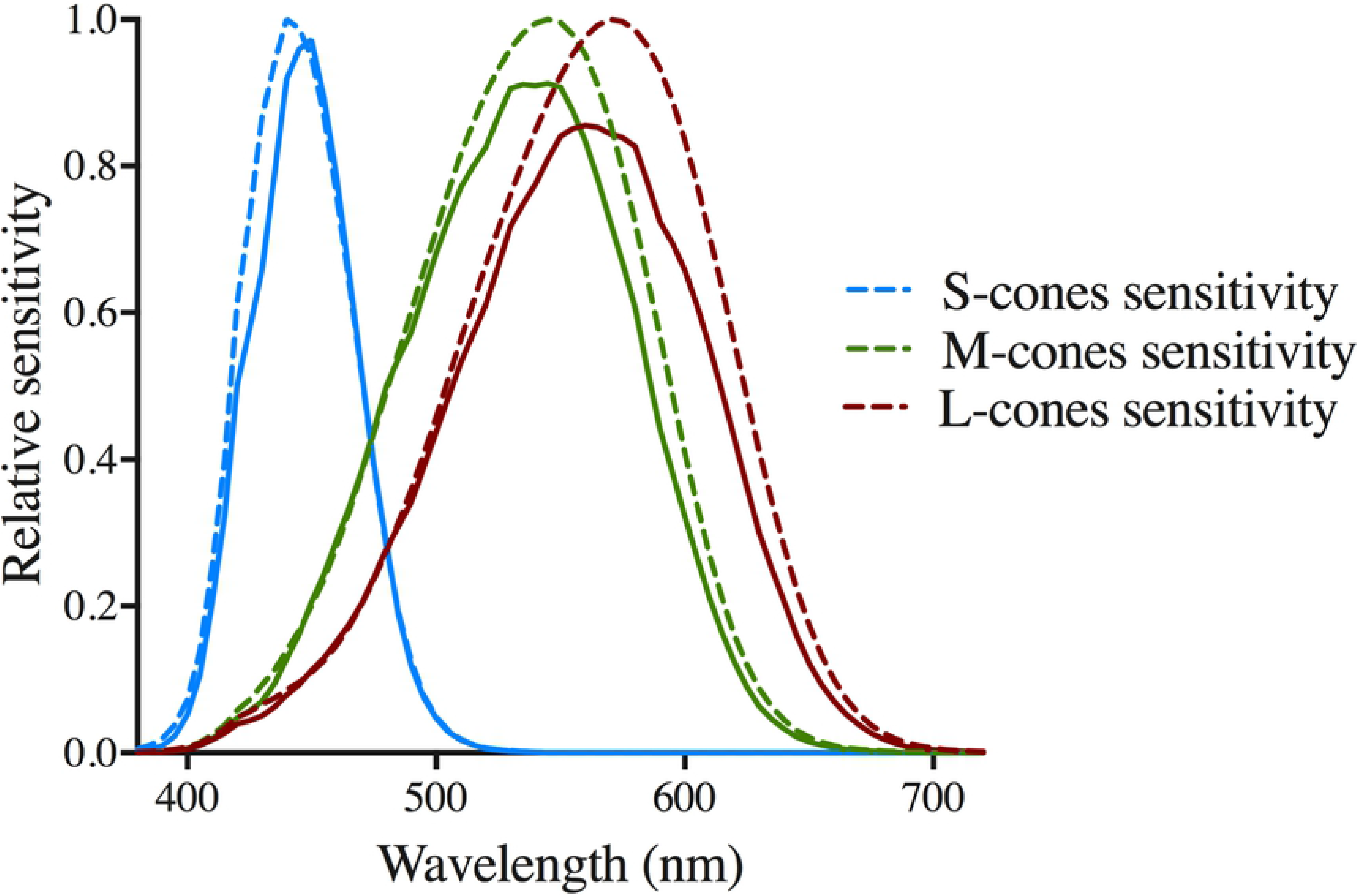
The relative sensitivities of S-cones. L-cones, and M-cones when are exposed to the LED lamp. Dashed lines represent the normal sensitivity of cone photoreceptors based on CIE TN 003:2015 [24], while solid lines represent the simulated sensitivity curves resulting from exposure to LED lamp.

The above mentioned procedures were repeated for achromatic targets with contrast levels of 0.1, 0.2 and 0.4 (corresponding to luminance values of 3.3, 3.6, and 4.2 cd/m^2^ and thus these stimuli were within the low phototopic range), and the four different BBL brands in a randomised order using an online tool (available at www.randomization.com). Each contrast level was repeated twice. Using the same procedures, testing was also conducted with chromatic red, green, blue and yellow stimuli set to a contrast of 0.4.

#### Results

The PSRTs required to detect an achromatic target stimulus were plotted against the stimulus luminance contrast (Fig 4). Different symbols represent different lens types, and error bars signify 1 standard error of the mean. A two-way repeated measures ANOVA showed a main effect of contrast such that increasing the stimulus contrast reduced PSRTs (F [2,18] = 4.013, p = 0.036). However, PSRTs between different BBLs and the control lens were not significantly different regardless of the contrast level (F [4,72] = 0.8, p = 0.5291). These results indicated that for achromatic stimuli presented under photopic conditions, while reducing contrast resulted in longer PSRTs, BBLs did not significantly affect PSRTs after photostress when compared to a clear control lens.

**Fig 4.**
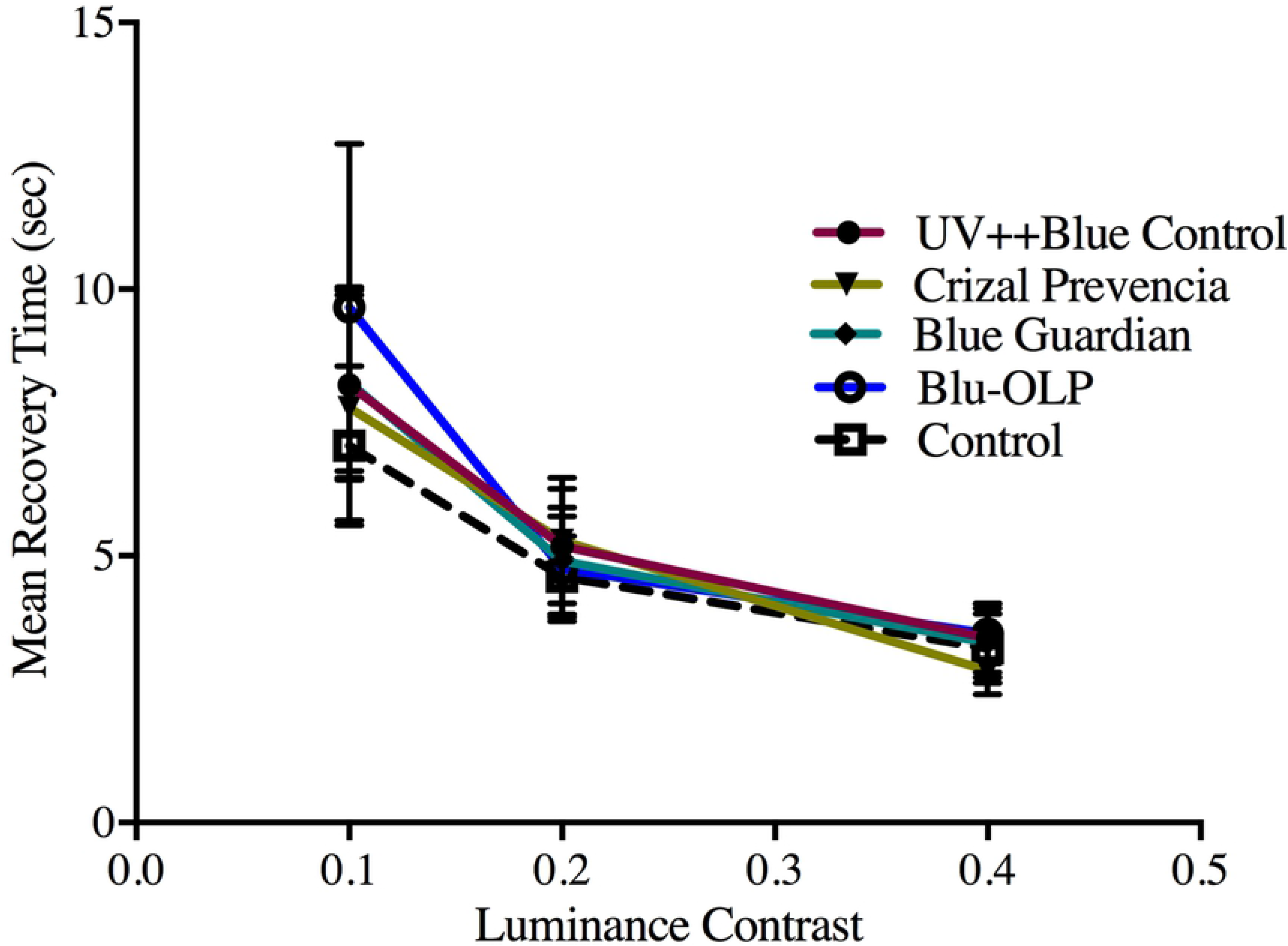
Mean recovery times are plotted as a function of luminance contrast for an achromatic stimulus. Error bars signify 1 SEM. All achromatic stimuli were high contrast and were tested for each lens type under photopic conditions of 3.3, 3.6, and 4.2 cd/m2.

The mean difference in PSRTs between each BBL type and the control lens required to detect different chromatic stimuli are shown in Fig 5. Error bars signify one standard error of the mean. A two-way repeated measures ANOVA (mixed effect) showed no significant effect of BBL type (F (9,72) = 0.490, p = 0.879), but there was a main effect of colour (F (3,72) = 4.29, p = 0.0156). These findings suggest that while the BBL type did not affect PSRTs, they were dependent on the stimulus colour. As evident in Fig 5, while PSRTs to red, green and yellow stimuli viewed through BBLs did not greatly differ from the control lens, however, blue stimuli resulted in a modest (approximately 1 – 2 seconds) increase in PSRTs. These results indicate that under photopic stimulus conditions, BBLs might have a small effect on PSRTs, particularly to a blue stimulus.

**Fig 5.**
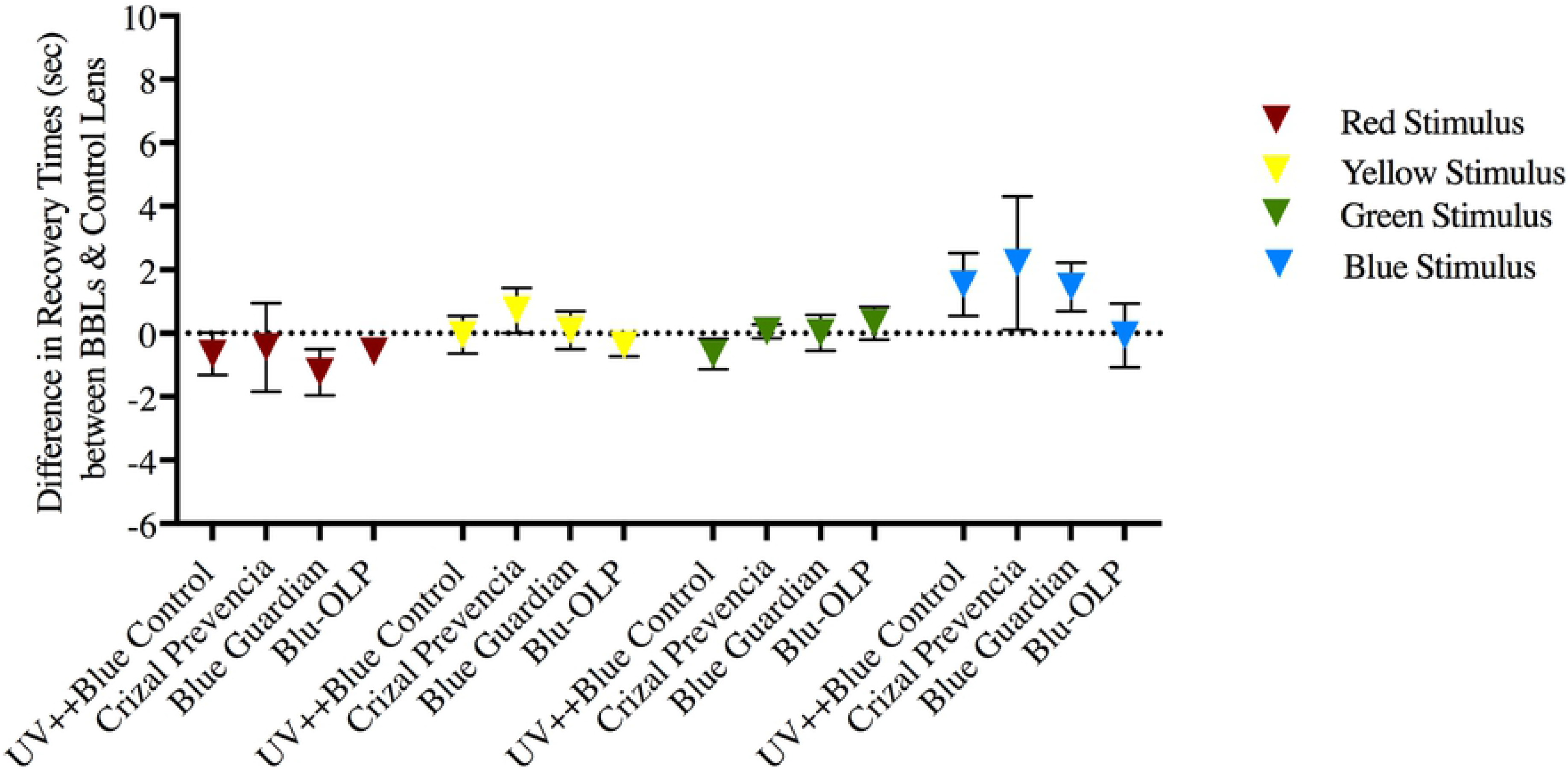
Mean difference in recovery times for high contrast coloured stimuli and lens types. Error bars signify one standard error of the mean. High contrast coloured stimuli can be red, yellow, green, or blue, and all stimuli were tested for each lens type under the photopic condition of 4.2 cd/m^2^.

### Experiment 2: Recovery times under mesopic conditions

In Experiment 2, we measured PSRTs at low luminance levels to simulate mesopic viewing conditions. As mentioned, previous studies investigating contrast sensitivity [5, 10] suggests that BBLs can greatly affect perception at low light-levels, and accordingly, it might be expected that PSRTs are longer as the overall light level available for perception. Moreover, the selective nature of BBLs might further render blue targets less perceptible and further increasing PSRTs. Here, we repeated Experiment 1 and a neutral density filter overlaid with the BBLs was used to reduce the overall light intensity by an approximately a factor of 10. Both achromatic and chromatic stimuli were tested at one contrast level of 0.4.

#### Results

Fig 6 shows the mean recovery time required to detect an achromatic target in Experiment 2. A one-way repeated measures ANOVA observed a significant effect of BBL type (F (2.006,8.02) = 61.95, p < 0.0001), indicating that PSRTs were dependent on the type of BBL. Post-hoc comparisons test (Dunnett’s test corrected for multiple comparisons assuming an alpha of 0.05) showed that both the Blu-OLP (mean difference, 6.475s, p = 0.0011) and Crizal Prevencia (mean difference, 3.038s, p = 0.0018) lenses produced significantly longer PSRTs than the control lens. However, PSRTs for the UV++Blue Control and Blue Guardian lenses were not significantly different from the control lens. Importantly, these results suggest that under low light levels, BBLs affect PSRTs, but this is dependent on the type of lens.

**Fig 6.**
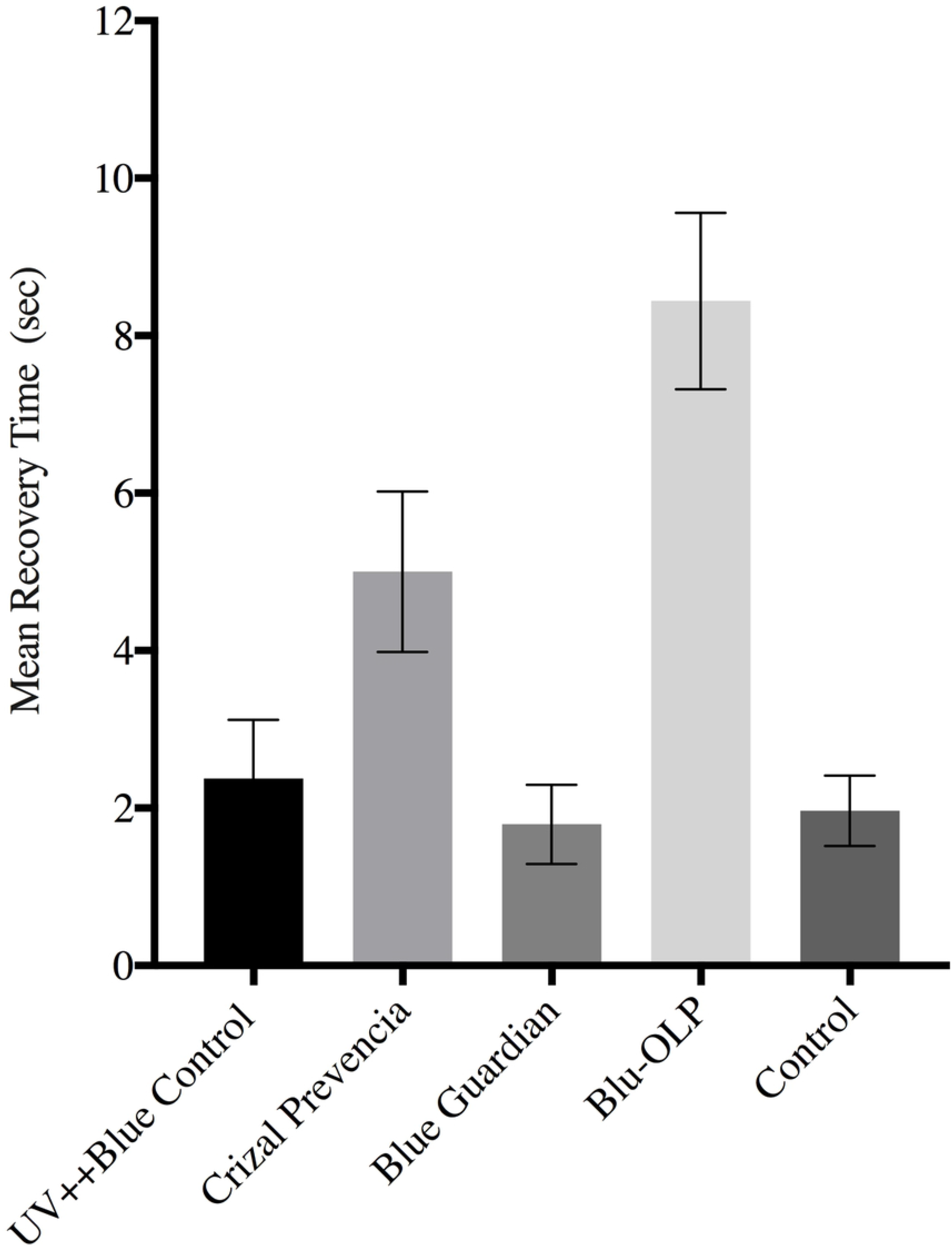
Mean recovery times for low contrast achromatic stimuli and lens types. Error bars signify one standard error of the mean. All achromatic stimuli were tested for each lens type under the mesopic condition of 0.4 cd/m2.

Note that the different lenses used in the present study differ in their transmission profiles (see Fig 1) and PSRTs might be dependent on the overall amount of light transmitted by these lenses. To investigate this, we calculated the area under the curve (AUC) values for the transmittance functions shown in Fig 1 and correlated them with their average PSRTs. Indeed, we find that PSRTs were significantly correlated with AUC with r = 0.964, p = 0.036. These results suggest that for achromatic stimuli BBLs affect PSRTs, but this is dependent on their overall light transmittance properties.

In Fig 7, the mean recovery time difference (relative to the clear control lens) are shown for different coloured stimuli (different plots) and BBL type. Error bars signify one standard error of the mean. A repeated-measures two-way ANOVA (mixed effect) was conducted with colour and BBL type as factors. This analysis observed a main effect of colour (F (3, 48) = 149.53, p < 0.0001) and BBL type (F (3,16) =139.01, p < 0.0001). A significant interaction effect was also evident (F (9, 48) = 278.79, p < 0.0001) which indicated that PSRTs for each colour were dependent on the type of BBL. Particularly, average PSRTs (relative to the control lens) for blue stimuli (38.40s) were considerably longer than yellow (0.7775s), green (0.1380s) and red (1.539s) stimuli. Indeed, Tukey’s multiple comparisons tests indicated that PSRTs to the blue stimulus was significantly longer than the other colours (Ps < 0.0001). One sample t-test showed that only the PSRTs for the blue stimulus (regardless of BBL type) was significantly different (t (19) = 5.726, p < 0.0001) from zero (i.e., the control lens). Given that BBLs selectively block short wavelengths, these results demonstrate that at low contrasts BBLs considerably affect the ability of the visual system to recover from photostress, which has implications for their wear under conditions of twilight and night time driving where overall light levels are low, and objects are frequently low contrast.

**Fig 7.**
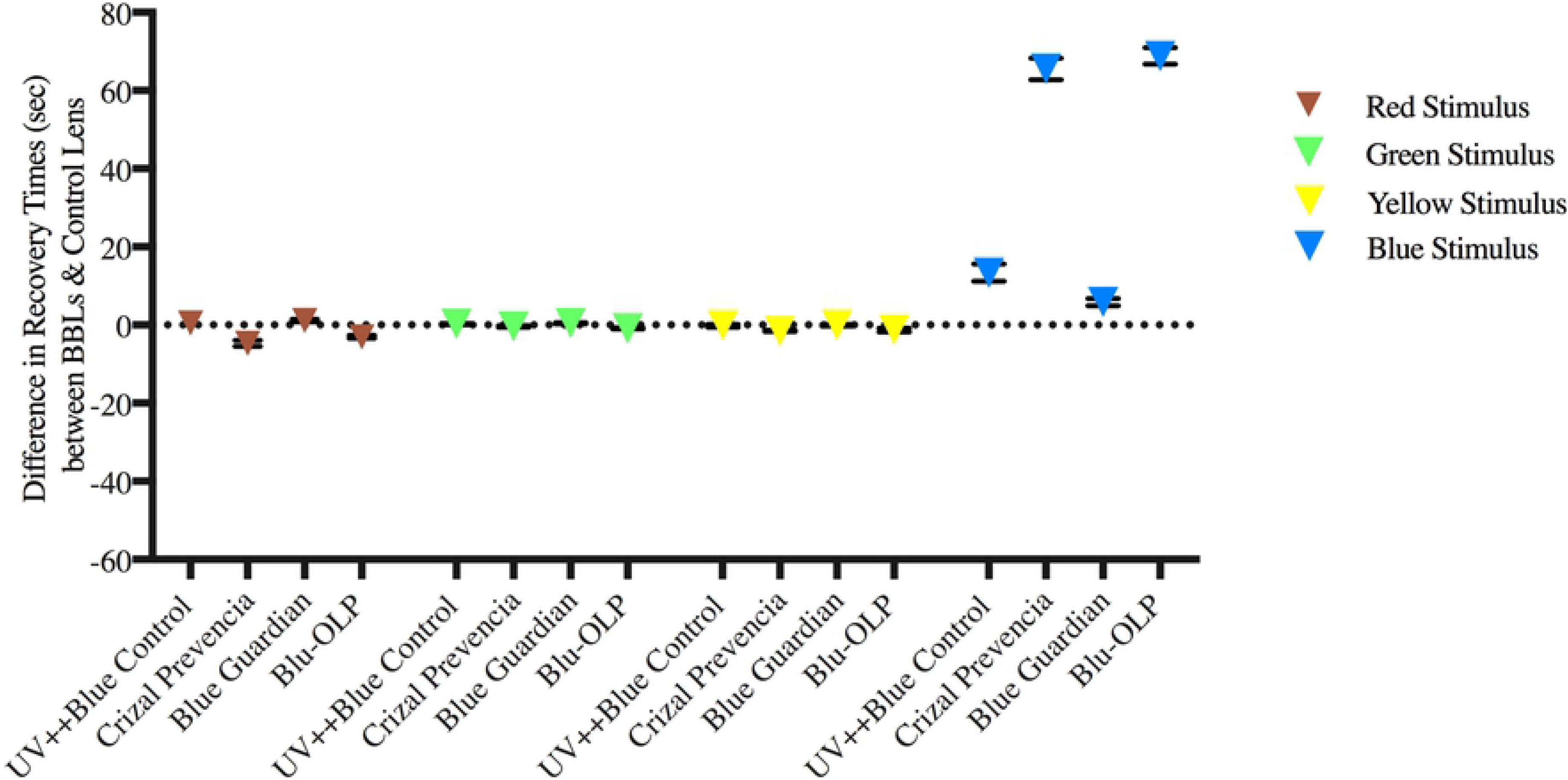
Mean difference in recovery times for low contrast coloured stimuli and lens types. Error bars signify one standard error of the mean. Low contrast coloured stimuli can be red, yellow, green, or blue, and all stimuli were tested for each lens type under the mesopic condition of 0.4 cd/m2.

A significant interaction effect indicated that the effect of BBLs on PSRTs was dependent on the colour of the stimulus. Given this, individual one-way repeated measures ANOVA was conducted for each coloured stimulus. This analysis showed that for red, green and yellow coloured targets, there was no significant difference between the BBLs, and their PSRTs were not significantly different from the clear lens. However, a significant effect was observed between BBLs for the blue coloured target (F (1.513, 6.050) = 155.3, p < 0.0001), and Post-hoc comparisons tests indicated that all BBLs led to significantly longer PSRTs relative to the control lens. In particular, the Crizal Prevencia and Blu-OLP resulted in much longer PSRTs (> 30 seconds) compared to the control lens. PSRTs are consistent with the transmittance properties of the BBLs, particularly the degree to which they block blue light. Indeed, a Spearman’s correlation between the transmittance efficiency for each BBL for the blue target (CIE 1931 xy: 0.15, 0.06) of a dominant wavelength of 440 nm used in the present study revealed a significant positive relationship (r = 0.994, p = 0.004).

## Discussion

In this study, the effect of wearing BBLs on PSRTs for low and high contrast stimuli was measured. We find that under photopic stimulus conditions, while reducing luminance contrast increased PSRTs, BBLs had a modest influence on PSRTs (relative to a clear control lens) for chromatic stimuli only (Fig 5). However, under mesopic stimulus conditions, BBLs significantly affect PSRTs for both achromatic and chromatic stimuli, particularly for blue coloured targets, which had considerably longer PSRTs. The type of BBL was also shown to selectively affect PSRTs, with those with transmittance profiles that block the most blue light having longer PSRTs.

The findings of our study demonstrate that the use of BBLs has unintended adverse consequences to visual function. Particularly, the fact that BBLs are designed to reduce light transmittance, albeit at shorter wavelengths, reduces the availability of light for visual perception. Indeed, some preliminary studies [5, 10] have shown impaired contrast sensitivity under low light conditions. Additionally, BBLs are known to induce a Tritan like defect in colour vision, and thus further impairing visual function [25, 26]. These findings, together with those reported by the present study, raise concerns and caution regarding their everyday use.

Our finding of longer PSRTs, particularly for blue stimuli under low light levels has important ramifications for night time activities or those in which light-levels are low. For example, during night time driving, the driver might be briefly exposed to bright lights by glancing at the headlights of a passing car. BBLs impair night time vision as they would unintendedly increase the time required for vision to be restored after bright light exposure. This might result in delayed object detection, and therefore stoppage and reaction times, and therefore might pose a safety risk for a driver, which outweighs their benefits, which has yet to be proven. These unintended properties of BBLs are particularly significant given that newer generation BBLs are a design feature of spectacles intended for everyday wear and cannot be removed.

In the present study, PSRTs were measured in younger participants (18-39 years old) with no history of ocular disease or abnormal vision. However, PSRTs have been shown to be dependent on age and are significantly longer due to eye disease. For older individuals, potential reductions in lens transparency, presence of vitreous floaters and slower response to light stimuli (particularly to coloured targets [27]) are common problems associated with ageing [28] and are likely to impact and increases PSRT. In addition, longer PSRT has been observed in individuals with primary open-angle glaucoma (POAG) and AMD [29–31], which are likely to further exacerbated by wearing BBLs. However, further research is needed to quantify the full extent to which BBL type affect vision in elderly people and patients with colour vision deficiency.

## Acknowledgments

The authors of this study wish to thank, Mr Justin Baker (JuzVision), Mr Tim Thurn (Essilor Australia) and Ms Dubravka Huber (Optometry Clinic, School of Optometry and Vision Science, UNSW Sydney) for providing BBLs used in this study. We thank Dr Kathleen Watt for valuable discussion.

